# Membrane Voltage Dynamics of Parvalbumin Interneurons Orchestrate Hippocampal Theta Rhythmicity

**DOI:** 10.1101/2022.11.14.516448

**Authors:** Hua-an Tseng, Rebecca A. Mount, Eric Lowet, Howard J. Gritton, Cyrus Cheung, Xue Han

**Affiliations:** Department of Biomedical Engineering, Boston University; Boston, Massachusetts, USA; Department of Comparative Biosciences, University of Illinois; Urbana, Illinois, USA

## Abstract

Hippocampal network activity at theta frequencies (5-10Hz) is important for behavior. However, it remains unclear how behaviorally-relevant network theta rhythms arise and interact with cellular dynamics to dictate spike timing. We performed membrane voltage (Vm) imaging of individual CA1 pyramidal cells and parvalbumin interneurons with simultaneous local field potential (LFP) recordings in mice during locomotion. We found that Vm theta rhythms organize spike timing in both cell types regardless of behavioral conditions, but the Vm of parvalbumin interneurons is better synchronized with LFP. The temporal relationships between spikes and LFP theta reliably reflect the Vm-LFP relationships in parvalbumin cells, but not in pyramidal cells. Thus, cellular theta rhythms broadly organize spike timing in CA1 neurons, and parvalbumin interneurons are critical in coordinating network theta rhythms.

**One-Sentence Summary:** Cellular membrane voltage of parvalbumin interneurons organizes spiking and network dynamics in the hippocampus.

## Introduction

Hippocampal rhythmic network activities, captured by local field potential (LFP) oscillations, have been broadly linked to learning and memory (*1-4*). Additionally, LFP oscillations have been shown to organize spike timing during behavior (*2, 5*). As spike occurrence is also influenced by cellular biophysical properties, anatomical connectivity patterns, and network states, different neuron subtypes exhibit variable temporal relationships with LFP oscillations. Because the hippocampus displays prominent LFP oscillations in the theta frequency range (5-10 Hz), many studies have characterized how spike timing in different neuron subtypes relate to LFP theta phase (*2, 5*). Interestingly, CA1 pyramidal cell spiking does not exhibit a consistent temporal relationship to LFP theta phase, likely due to variations in behavioral conditions (*1, 6-9*). A well-known example is theta phase precession, in which the spiking of a CA1 place cell gradually shifts to earlier phases of LFP theta as the animal traverses the place field of that particular cell (*3, 4, 10*). In contrast, hippocampal interneurons, particularly parvalbumin-expressing (PV) interneurons, exhibit much more consistent spike-LFP theta phase relationships (*11-13*). However, the cellular mechanisms linking LFP theta oscillations and spike timing during behavior remain elusive.

PV cells play an important role in supporting CA1 LFP theta oscillations. Optogenetic activation or suppression of PV cells respectively enhances or reduces CA1 LFP theta oscillations in hippocampal brain slices (*14*), and genetically ablating synaptic inhibition onto PV interneurons *in vivo* reduces the CA1 LFP theta oscillation (*15*). Further, CA1 PV interneurons are the primary target of the GABAergic medial septum, which exhibits theta frequency rhythmic activity (*2, 16-20*). Thus, PV cells are hypothesized to relay rhythmic septal input, thus entraining downstream neurons, including pyramidal cells and other interneurons, via direct inhibitory synaptic input. Indeed, rhythmic depolarization of GABAergic basket cells and axo-axonic cells (both largely PV cells) at theta frequencies can synchronize both spiking and subthreshold voltage dynamics of CA1 pyramidal cells (*21*). Similarly, rhythmic optogenetic activation of PV cells at theta frequencies paces pyramidal cell spiking *in vivo* (*7*). Finally, inhibitory inputs from PV cells can modulate the relationship between pyramidal cell spike timing and LFP theta phase (*1, 6*). These results support the critical role of PV cells in organizing pyramidal cell spiking to LFP theta phases.

Not only are theta oscillations prominent at the network level of the CA1, but intracellular studies have demonstrated that the membrane potential (Vm) of individual hippocampal neurons shows theta rhythmicity as well (*22-27*). Vm is shaped by both synaptic inputs and the biophysical and morphological properties of the neuron (*1, 28*). Since spike generation relies on Vm depolarization, Vm oscillations of individual neurons provide a critical cellular link between network LFP oscillations and the spiking output of individual cells within that network. However, due to the technical difficulty of performing intracellular electrophysiological recordings in specific neuron types in behaving mammals, there has been limited *in vivo* evidence on how Vm relates to LFP and spike timing (*24, 25, 29-33*).

To investigate how cellular Vm oscillations of individual neurons link LFP theta oscillations and spike timing, we performed *in vivo* voltage imaging of individual CA1 pyramidal cells and PV cells in awake animals during resting and walking. These behavioral states induce different strengths of LFP theta power in the CA1. We targeted CA1 pyramidal and PV interneurons through cell-type-specific expression of SomArchon, a high-performance genetically-encoded voltage sensor that reports Vm at the soma (*26, 34*). We characterized the relationships between Vm theta oscillations, spike timing, and LFP theta oscillations in PV interneurons versus pyramidal cells during the two locomotor states.

## Results

### Both CaMKII-positive pyramidal cells and PV interneurons exhibit prominent membrane voltage (Vm) theta oscillations across behavioral states with low and high LFP theta power

To investigate how membrane potential (Vm) of dorsal CA1 pyramidal cells versus PV-positive interneurons relates to CA1 LFP, we performed simultaneous voltage imaging and LFP recording through a chronically-implanted imaging window above the pyramidal cell layer coupled to an electrode ∼200µm below the imaging area (Figure 1A, E, and F, also see Methods). AAV9-CaMKII-SomArchon-GFP was used to express SomArchon in CaMKII-positive neurons that are primarily pyramidal cells (CaMKII cells; N=31 neurons from 4 mice, all with simultaneous LFP). AAV9-FLEX-SomArchon-GFP was used in PV-Cre mice to express SomArchon specifically in PV-positive cells (PV cells; N=48 neurons from 9 mice, 28 cells with simultaneous LFP) (Supplemental Figure 1). During each recording session, SomArchon voltage imaging and LFP recording were performed while mice were resting or walking at a rate of 11.75 cm/s on a motorized treadmill (Figure 1A).

**Figure 1.**
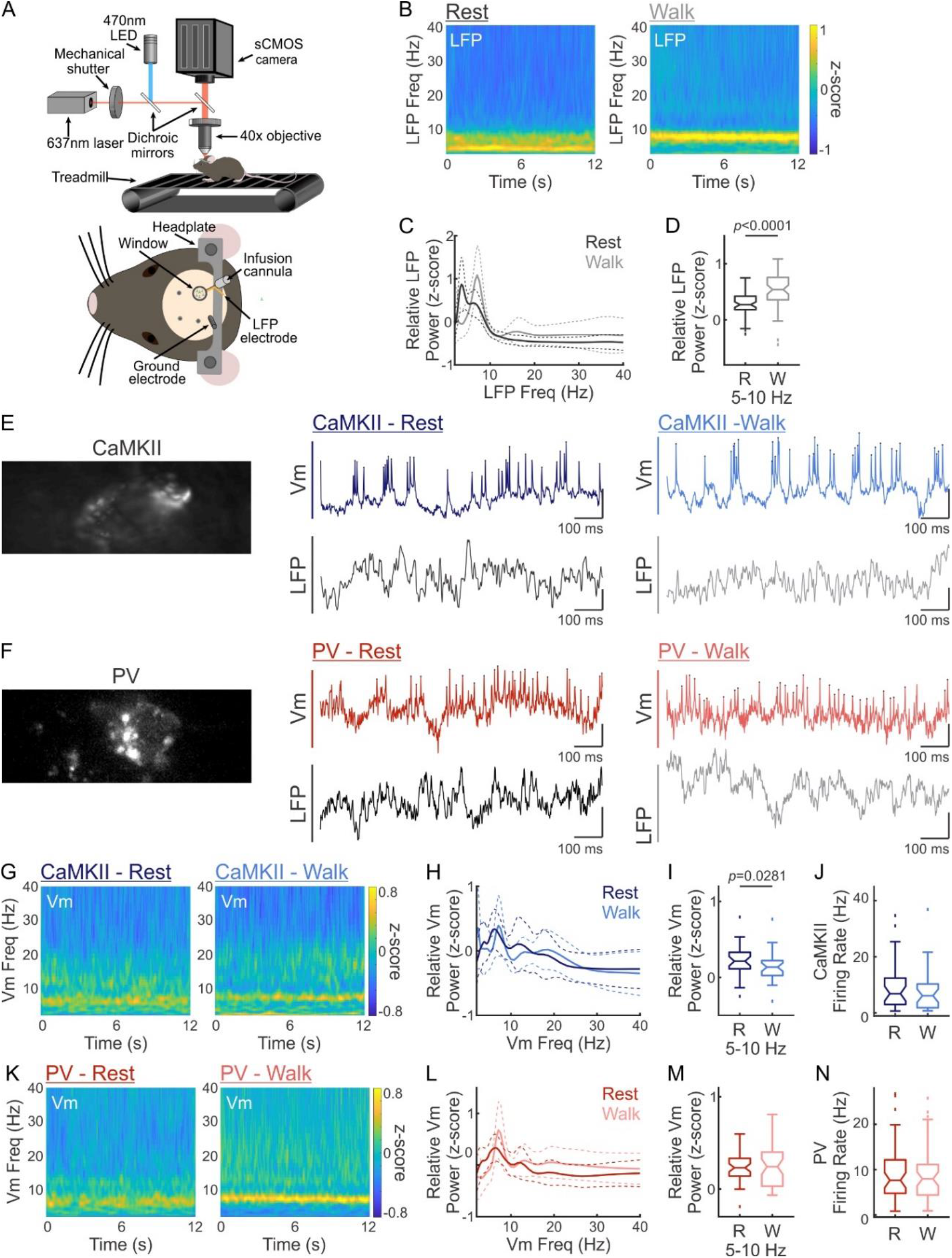
Membrane voltage (Vm) dynamics of CaMKII-positive pyramidal neurons and PV-positive interneurons during resting and during locomotion. (**A**) Experimental setup (top) and imaging apparatus design (bottom). During imaging, mice alternated between resting and walking. The imaging head implant consists of a glass window coupled to a LFP recording electrode. (**B**) Mean LFP spectrograms from all recordings during resting (left) and walking (right) (N=59 cells). (**C**) LFP power distributions of all recordings during resting (black) and walking (gray) (N=59 cells). (**D**) LFP theta power during resting (black) and walking (gray) (N=59 cells, Wilcoxon signed rank test, p<0.0001, resting: 0.29±0.20, running: 0.55±0.30). (**E**) Mean SomArchon intensity of an example CaMKII neuron (left), its optically recorded membrane potential (Vm, SomArchon signal), and simultaneously recorded LFP during resting (middle, Vm: dark blue, LFP: black) and running (right, Vm: light blue, LFP: gray). (**F**) Mean SomArchon intensity of an example PV neuron (left), its Vm, and simultaneously recorded LFP during resting (middle, Vm: dark red, LFP: black) and running (right, Vm: light red, LFP: gray). (**G**) Mean Vm spectrograms from all CaMKII neurons during resting (left) and walking (right) (N=31 cells). (**H**) Mean Vm power distributions of the CaMKII population during resting (dark blue) and walking (light blue) (N=31 cells). (**I**) CaMKII Vm theta power during resting (dark blue) and walking (light blue) (N=31 cells, Wilcoxon signed rank test, p=0.0282, resting: 0.22±0.21, walking: 0.14±0.20). (**J**) Firing rate of CaMKII cells during resting (dark blue) and walking (light blue) (N=31 cells, Wilcoxon signed rank test, p=0.0938, resting: 9.73±8.84Hz, walking: 7.74±7.76Hz). (**K**) Mean Vm spectrograms from all PV neurons during resting (left) and walking (right) (N=48 cells). (**L**) Vm power distributions of the PV population during resting (dark red) and walking (light red) (N=48 cells). (**M**) Mean PV Vm theta power during resting (dark red) and walking (light red) (N=48 cells, Wilcoxon signed rank test, p=0.2680, resting: 0.23±0.15, walking: 0.27±0.24). (**N**) Firing rate of PV cells during resting (dark red) and walking (light red) (N=48 cells, Wilcoxon signed rank test, p=0.5485, resting: 9.36±6.48Hz, walking: 8.54±5.76Hz). In power distribution plots, the solid lines and the dashed lines indicate mean and ±standard deviation, respectively. The power at each time point was normalized by calculating its z-score across all frequencies. In boxplots, notch indicates median, box indicates 25th to 75th percentiles, whiskers indicate the range of non-outliers, and dots indicate outliers.

Since hippocampal LFP theta oscillations have been broadly documented during locomotion (*2, 35-38*), we first compared LFP oscillation power between resting and walking and found a significant increase in LFP theta power during walking (Figure 1B-D, N=59 cells, 31 CaMKII cells and 28 PV cells, Wilcoxon signed rank test, p<0.0001, resting: 0.29±0.20, running: 0.55±0.30, also see Supplemental Figure 2). LFP measures extracellular field potentials around the electrode site, capturing aggregate currents across the somatic, dendritic, and axonal membranes of the local brain area. To examine how these network-level CA1 LFP oscillations relate to cellular-level Vm dynamics in individual CA1 neurons, we recorded somatic transmembrane voltage-dependent SomArchon fluorescence (Vm) with a wide-field microscope at 827Hz.

We detected prominent Vm theta oscillations in both CaMKII cells and PV interneurons during both resting and walking (Figure 1G and K, respectively). Despite the increase in LFP theta power during walking, CaMKII Vm theta power decreased during walking compared to resting (Figure 1H and I, N=31 cells, Wilcoxon signed rank test, p=0.0282, resting: 0.22±0.21, walking: 0.14±0.20) and PV Vm theta power remained constant between the two conditions (Figure 1L and M, N=48 cells, Wilcoxon signed rank test, p=0.2680, resting: 0.23±0.15, walking: 0.27±0.24). Thus, Vm theta oscillations are prominent cellular features of both CaMKII and PV neurons, and cellular theta oscillation power is not enhanced by walking, unlike network LFP theta power.

### Spike timing in both CaMKII and PV neurons is organized by Vm theta oscillation phase

Having established that both CaMKII and PV neurons exhibited prominent Vm theta oscillations, we then evaluated how spikes in CaMKII and PV neurons relate to Vm theta oscillations. We first identified spikes in the recorded Vm traces as previously described (see Methods). We found no significant differences in the firing rates of either population between the two behavioral conditions (Figure 1J and N, CaMKII cells: N=31 cells, Wilcoxon signed rank test, p=0.0938, resting: 9.73±8.84Hz, walking: 7.74±7.76Hz, PV cells: N=48 cells, Wilcoxon signed rank test, p=0.5485, resting: 9.36±6.48Hz, walking: 8.54±5.76Hz).

Oscillations can be characterized by the temporal component (oscillation phase) and the amplitude component (oscillation power), and here we first considered phase. Previous intracellular patch clamp recording and voltage imaging studies in CA1 neurons have shown that spikes are generally more phase-locked to Vm theta oscillations than LFP theta oscillations (*24-26*). However, it is unknown whether spike-Vm theta phase-locking in individual neurons is modulated across behavioral states with enhanced or reduced LFP theta power. We found that most CaMKII and PV cells showed significant spike-Vm theta phase-locking regardless of behavioral condition, with similar fractions of phase-locked cells in each population (CaMKII: Figure 2A, N=31 cells, resting: 77%, walking: 71%, Fisher’s exact test, p=0.7723; PV: Figure 2C, N=48 cells, resting: 83%, walking: 81%, Fisher’s exact test, p=1.0000). Additionally, the strength of spike-Vm theta phase-locking across neurons was also similar during both resting and walking (CaMKII: Figure 2B, N=31 cells, Wilcoxon signed rank test, p=0.8909, resting: 0.37±0.15, walking: 0.36±0.11, PV: Figure 2D, N=48 cells, Wilcoxon signed rank test, p=0.3560, resting: 0.38±0.13, walking: 0.36±0.13).

**Figure 2.**
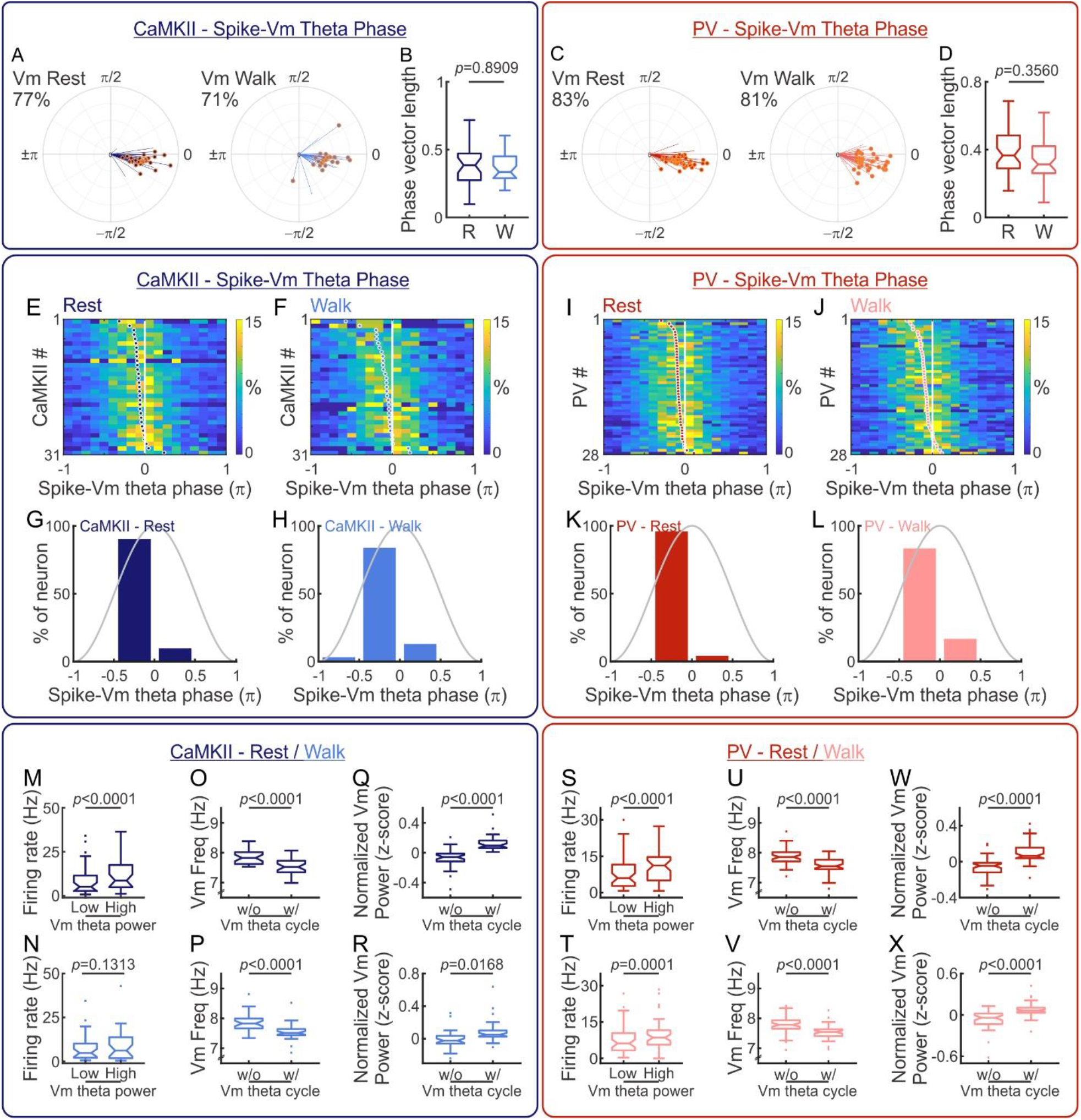
Vm theta oscillations organize spike timing in both cell types. (**A**) Average spike phase relative to the Vm theta oscillation of individual CaMKII neurons during resting (left) and walking (right) (N=31 cells). The orange circle at the end of the phase vector indicates significant phase-locking. The number indicates the percentage of neurons showing significant phase-locking. (**B**) Amplitude of spike phase-locking relative to Vm theta oscillation in the CaMKII population during resting (dark blue) and walking (light blue) (N=31, Wilcoxon signed rank test, p=0.8909, resting: 0.37±0.15, walking: 0.36±0.11). (**C**) Average spike phase relative to Vm theta oscillation of individual PV neurons during resting (left) and walking (right) (N=48 cells). The orange circle at the end of the phase vector indicates significant phase-locking. The number indicates the percentage of neurons showing significant phase-locking. (**D**) Amplitude of spike phase-locking relative to Vm theta oscillation in the PV population during resting (dark red) and walking (light red) (N=48, Wilcoxon signed rank test, p=0.3560, resting: 0.38±0.13, walking: 0.36±0.13). (**E** and **F**) Distributions of spike phase relative to the Vm theta oscillation for individual CaMKII neurons during resting (E) and walking (F). Each row shows the spike phase distribution of one neuron, and neurons were sorted by their average spike phase, indicated by the dots. (**G** and **H**) Histograms of average spike phase relative to Vm theta oscillation for all CaMKII neurons during resting (G) and walking (H) (N=31 cells). (**I** and **J**) Distribution of spike phase relative to the Vm theta oscillation for individual PV neurons during resting (I) and walking (J). Each row shows the spike phase distribution of one neuron, and neurons were sorted by their average spike phase, indicated by the dots. (**K** and **L**) Histograms of average spike phase relative to Vm theta oscillation for all PV neurons during resting (K) and walking (L) (N=28 cells). (**M** and **N**) CaMKII firing rates during Vm theta cycles with low and high Vm theta power during resting (M, N=31 cells, Wilcoxon signed rank test, p=7.202e-6, low power firing rate: 8.54±8.79Hz, high power firing rate: 12.11±9.58Hz) and walking (N, N=31 cells, Wilcoxon signed rank test, p=0.1313, low power firing rate: 7.72±7.62Hz, high power firing rate: 8.66±8.74Hz). (**O** and **P**) Peak frequency of CaMKII Vm theta cycles with and without spikes, during resting (O, N=31 cells, Wilcoxon signed rank test, p=1.578e-6, without spike frequency: 7.84±0.24Hz, with spike frequency: 7.53±0.27Hz) and walking (P, N=31 cells, Wilcoxon signed rank test, p=2.114e-5, without spike frequency: 7.87±0.33Hz, with spike frequency: 7.55±0.28Hz). (**Q** and **R**) Normalized theta power of CaMKII Vm theta cycles with and without spikes, during resting (Q, N=31 cells, Wilcoxon signed rank test, p=9.475e-6, without spike power: -0.07±0.13, with spike power: 0.12±0.10) and walking (R, N=31 cells, Wilcoxon signed rank test, p=0.0168, without spike power: -0.01±0.11, with spike power: 0.08±0.14). (**S** and **T**) PV firing rates during Vm theta cycles with low and high Vm theta power during resting (S, N=48 cells, Wilcoxon signed rank test, p=6.712e-6, low power firing rate: 8.13±6.98Hz, high power firing rate: 10.89±6.23Hz) and walking (T, N=48 cells, Wilcoxon signed rank test, p=0.0001, walking-low power firing rate: 7.68±5.80Hz, resting-high power firing rate: 9.63±6.25Hz). (**U** and **V**) Peak frequency of PV Vm theta cycles with and without spikes, during resting (U, N=48 cells, Wilcoxon signed rank test, p=1.111e-8, without spike frequency: 7.86±0.28Hz, with spike frequency: 7.56±0.27Hz) and walking (V, N=48 cells, Wilcoxon signed rank test, p=6.900e-7, without spike frequency: 7.78±0.27Hz, with spike frequency: 7.54±0.23Hz). (**W** and **X**) Normalized theta power of PV Vm theta cycles with and without spikes, during resting (W, N=48 cells, Wilcoxon signed rank test, p=1.431e-6, without spike power: -0.06±0.10, with spike power: 0.11±0.12) and walking (X, N=48 cells, Wilcoxon signed rank test, p=5.529e-6, without spike power: -0.06±0.13, with spike power: 0.07±0.10). In boxplots, notch indicates median, box indicates 25th to 75th percentiles, whiskers indicate the range of non-outliers, and dots indicate outliers.

In both CaMKII and PV neurons, almost all spikes occurred on the rising phase of the Vm theta cycle, during Vm depolarization, regardless of behavioral condition (CaMKII: Figure 2E and F, PV: Figure 2I and J). For both cell types, over 80% of neurons showed average spike-Vm theta phase-locking on the second half of the rising phase, with spikes occurring just before the peak of Vm theta oscillation (CaMKII: Figure 2G and H, PV: Figure 2K and L). These results show that spike timing is consistently coupled to the rising phase of the Vm theta oscillation in both pyramidal cells and PV interneurons, and this phase relationship is insensitive to behavioral conditions. The preferred occurrence of spikes on the rising phase provides direct evidence in behaving animals that spike generation threshold is not restricted to a fixed voltage. Instead, spikes preferentially occur during the monotonic depolarization of the rising phase, rather than the monotonic repolarization of the falling phase, of a Vm theta cycle, consistent with previous studies (*39-43*).

### Spike occurrence is associated with prolonged Vm theta cycles and elevated Vm theta power

We next examined how spike occurrence influence Vm theta oscillations. Vm theta power fluctuates during behavior; thus, to probe the relationship between spiking and Vm theta power, we calculated instantaneous theta power for each individual theta cycle and split the cycles into those with high (>0.5) versus low (<-0.5) power. In CaMKII cells, high-power Vm theta cycles were associated with higher firing rates than low-power cycles during resting, but not walking (Figure 2M and N, respectively, N=31 cells, Wilcoxon signed rank test, resting p=7.202e-6, resting-low power firing rate: 8.54±8.79Hz, resting-high power firing rate: 12.11±9.58Hz, walking p=0.1313, walking-low power firing rate: 7.72±7.62Hz, resting-high power firing rate: 8.66±8.74Hz). In contrast, in PV cells, high-power Vm theta cycles were associated with higher firing rates regardless of behavioral condition (Figure 2S and T, respectively, N=48 cells, Wilcoxon signed rank test, resting p=6.712e-6, resting-low power firing rate: 8.13±6.98Hz, resting-high power firing rate: 10.89±6.23Hz, walking p=0.0001, walking-low power firing rate: 7.68±5.80Hz, resting-high power firing rate: 9.63±6.25Hz). Overall, these results suggest that PV Vm theta power reliably determines spike generation, whereas the relationship between CaMKII Vm theta power and firing rate is behaviorally modulated.

*In vitro* electrophysiology studies have also widely documented that the occurrence of spikes can alter ongoing Vm dynamics by changing voltage-dependent conductance across the soma and dendrites (*44-46*). For example, spiking in hippocampal pyramidal cells is sometimes followed by prolonged depolarization, known as after-depolarization potential, that can last for tens of milliseconds (*47*). Therefore, we next explored whether spiking influences the length of Vm theta cycle and the amplitude of Vm theta power. Indeed, we found that Vm theta cycles that included spikes were significantly longer and had higher theta power than cycles without spikes in both cell types, under both behavioral conditions (CaMKII: Figure 2O-R, N=31 cells, Wilcoxon signed rank test, see figure for p-values, PV: Figure 3U-X, N=48 cells, Wilcoxon signed rank test, see figure for p-values). Thus, the occurrence of spikes is associated with prolonged Vm theta cycles and elevated Vm power, indicating a close relationship between spikes and Vm theta oscillations regardless of cell type or behaviorally-evoked network state.

**Figure 3.**
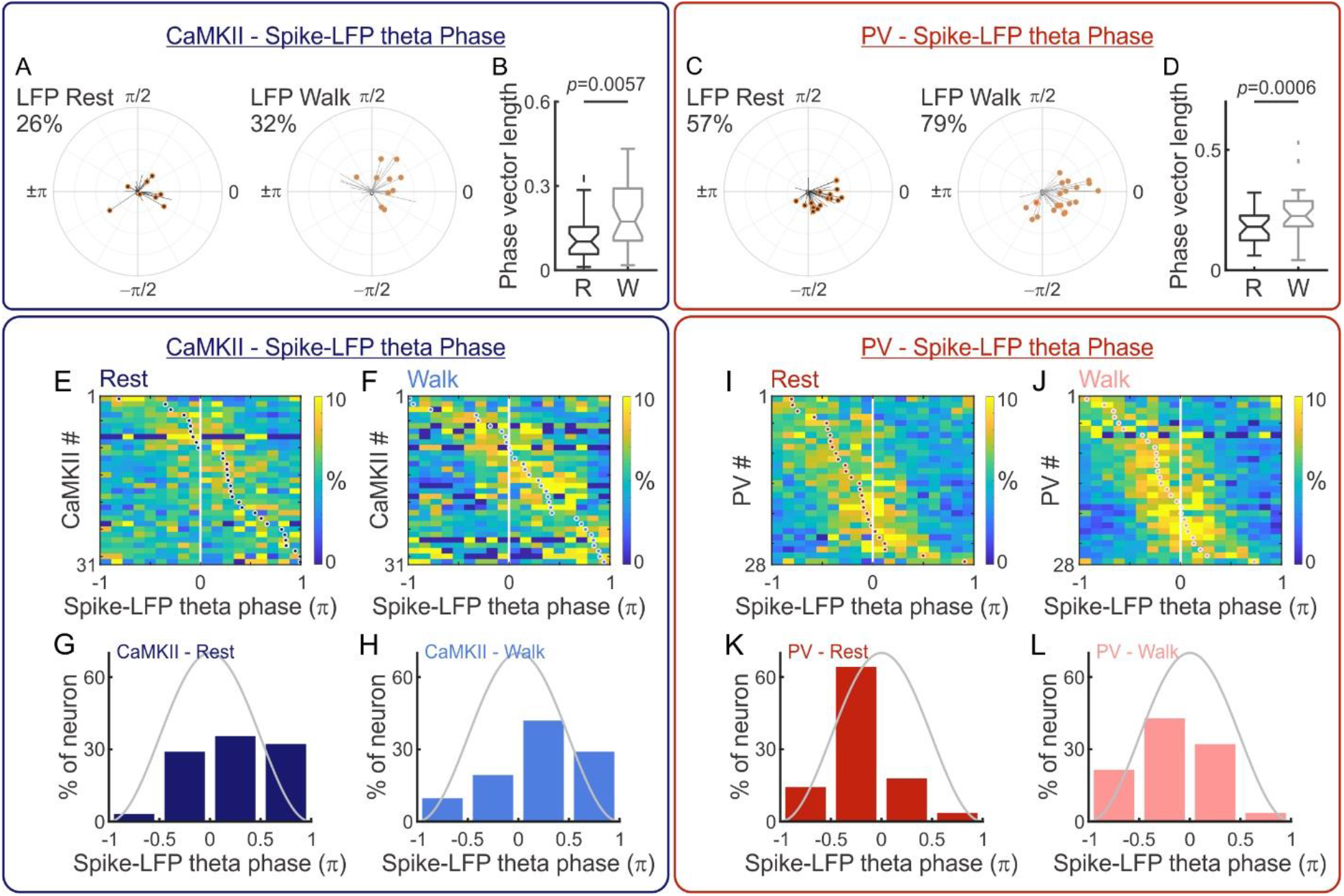
Distinct LFP phase preference of spikes in CaMKII versus PV neurons. (**A**) Average spike phase relative to LFP theta oscillation of individual CaMKII neurons during resting (left) and walking (right) (N=31 cells). The orange circle at the end of the phase vector indicates significant phase-locking. The number indicates the percentage of neurons showing significant phase-locking. (**B**) Amplitude of spike phase-locking relative to LFP theta oscillation across the CaMKII population during resting (black) and walking (gray) (N=31 cells, Wilcoxon signed rank test, p=0.0057, resting: 0.12±0.09, walking: 0.19±0.12). (**C**) Average spike phase relative to the LFP theta oscillation of individual PV neurons during resting (left) and walking (right) (N=28 cells). The orange circle at the end of the phase vector indicates significant phase-locking. The number indicates the percentage of neurons showing significant phase-locking. (**D**) Amplitude of spike phase-locking relative to the LFP theta oscillation of the PV population during resting (black) and walking (gray) (N=28 cells, Wilcoxon signed rank test, p=0.0006, resting: 0.18±0.07, walking: 0.24±0.11). (**E** and **F**) Distributions of spike phase relative to LFP theta oscillation for individual CaMKII neurons during resting (E) and walking (F). Each row shows the spike phase distribution of one neuron and the neurons were sorted by their average spike phase, indicated by the dots. (**G** and **H**) Histograms of average spike phase relative to LFP theta oscillation for all CaMKII neurons during resting (G) and walking (H) (N=31 cells). (**I** and **J**) Distributions of spike phase relative to LFP theta oscillation for individual PV neurons during resting (I) and walking (J). Each row shows the spike phase distribution of one neuron, and neurons were sorted by their average spike phase, indicated by the dots. (**K** and **L**) Histograms of average spike phase relative to LFP theta oscillation for all PV neurons during resting (K) and walking (L) (N=28 cells).

### PV neurons show more consistent spike-LFP theta phase-locking than CaMKII neurons

Like Vm oscillations, LFP oscillations can be described by both phase and power components. We first focused on the relationship between spikes and LFP theta phase. Previous extracellular recording studies demonstrated that CA1 pyramidal cell spiking exhibits diverse temporal relationships to LFP theta phase (*1, 3, 4, 6-10*). In contrast, spikes in fast-spiking interneurons (putative PV cells) generally occur at a narrow range of LFP theta phase (*11, 12*). As we detected a prominent increase in LFP theta power during walking (Figure 1), we asked whether spike-LFP phase relationships were modulated by behavioral state, even though the spike-Vm phase relationships we observed were insensitive to behavioral changes. We found that the strength of spike-LFP theta phase-locking significantly increased during walking for both CaMKII and PV populations compared to resting (CaMKII: Figure 3A and B, N=31 cells, Wilcoxon signed rank test, p=0.0057, resting: 0.12±0.09, walking: 0.19±0.12, PV: Figure 3C and D, N=28 cells, Wilcoxon signed rank test, p=0.0006, resting: 0.18±0.07, walking: 0.24±0.11). The fraction of cells that exhibited significant spike-LFP theta phase-locking did not change between behavioral conditions for either cell type (CaMKII: Figure 3A, N=31 cells, resting: 26%, walking: 32%, Fisher’s exact test, p=0.7802, PV: Figure 3C, N=28 cells, resting: 57%, walking: 79%, Fisher’s exact test, p=0.1516). However, a larger fraction of PV cells exhibited spike-LFP theta phase-locking compared to CaMKII cells during both behavioral conditions (Figure 3A and C, Fisher’s exact test, resting: p=0.0185, walking: p=0.0006).

To further examine spike-LFP phase relationships at the individual neuron level, we calculated a probability distribution of spikes relative to LFP theta phase for each neuron (Figure 3 E, F, I, and J). Across the entire CaMKII population, the spike-LFP theta phase of individual neurons spread throughout LFP theta phase regardless of behavioral condition (Figure 3G and H), consistent with previous work showing diverse phase-locking of pyramidal cell spiking to LFP theta (*1, 6-9*). In contrast, spike-LFP phase locking for individual PV cells concentrated around the late rising phase of LFP theta during resting and became more heterogeneous across the entire rising phase during walking (Figure 3K and L). The spike-LFP phase distributions of CaMKII and PV cells were significantly different from one another during both behavioral conditions (Chi-squared test, resting: p<0.0001, walking: p=0.0012).

We then investigated the power component of LFP theta relative to spikes, and found that LFP power was not coupled to spike timing under either locomotion condition in either cell type (Supplemental Figure 3). The lack of change in LFP power around spikes is consistent with the idea that there are broad and heterogeneous sources contributing to LFP signal, and thus individual neuron spiking alone is not sufficient to generate significant LFP oscillations in CA1 (*5*).

### PV Vm theta oscillations are more synchronized with LFP theta oscillations and the synchronization of PV Vm and LFP is accompanied by elevated LFP theta power

Because of the unique relationships observed between spiking and cellular-level Vm theta versus network-level LFP theta oscillations, we next assessed the phase relationship of Vm and LFP. Specifically, we calculated the phase shifts between individual Vm theta cycles and LFP theta cycles. We found that the phase shifts between CaMKII Vm theta and LFP theta were highly diverse, ranging from –π to π, during both resting and walking (Figure 4A-D), demonstrating that CaMKII Vm theta is generally not synchronized with LFP theta. In contrast, the phase shifts between PV Vm theta and LFP theta were concentrated around zero regardless of the animal’s behavioral state, indicating a high degree of synchronization between PV Vm theta and LFP theta (Figure 4G-I). In both behavioral states, the distributions of Vm-LFP theta phase shifts were significantly different between the CaMKII and PV populations (Chi-squared test, resting: p=0.0018, walking: p=0.0005), and the temporal deviations between Vm theta and LFP theta were significantly smaller in PV cells than in CaMKII neurons (Supplemental Figure 4, rank sum test, resting: p=0.0383, walking: p=0.0026). These observations indicate that Vm theta oscillations in PV neurons are more synchronized with LFP theta than those in CaMKII cells, regardless of behavioral condition.

**Figure 4.**
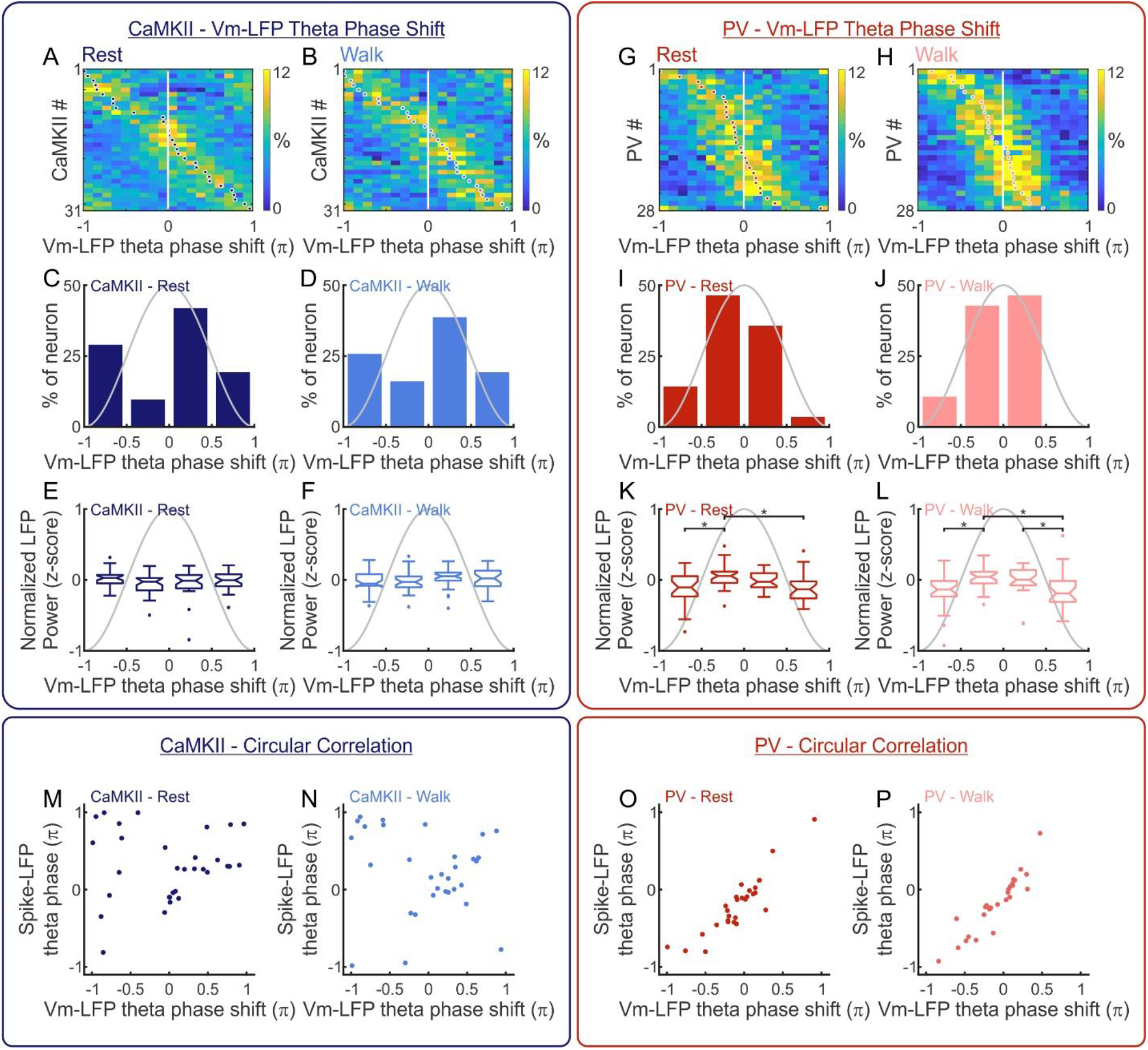
Distinct synchrony between Vm and LFP oscillations in CaMKII and PV neurons. (**A** and **B**) Distributions of the phase shifts between Vm theta cycles and LFP theta cycles of individual CaMKII neurons during resting (A) and walking (B). Each row shows the phase shift distribution of one neuron, and neurons were sorted by their average phase shift, indicated by the dots. (**C** and **D**) Histograms of average Vm-LFP phase shift for all CaMKII neurons during resting (C) and walking (D) (N=31 cells). (**E** and **F**) Normalized LFP theta power during the CaMKII Vm theta cycles with various phase shifts during resting (E) and during walking (F) (N=31 cells, one-way ANOVA). (**G** and **H**) Distributions of the phase shifts between Vm theta cycles and LFP theta cycles of individual PV neurons during resting (G) and walking (H). Each row shows the phase shift distribution of one neuron and the neurons were sorted by their average phase shift, indicated by the dots. (**I** and **J**) Histograms of average Vm-LFP phase shift for all PV neurons during resting (I) and walking (J) (N=28 cells). (**K** and **L**) Normalized LFP theta power during the PV Vm theta cycles with various phase shifts during resting (K) and walking (L) (N=28 cells, one-way ANOVA, *p<0.05). (**M** and **N**) Circular correlations between the average Vm-LFP theta phase shift and the average spike phase relative to the LFP theta oscillation for all CaMKII neurons during resting (M, N=31 cells, circular correlation coefficient=0.5313) and during walking (N, N=31 cells, circular correlation coefficient=0.5257). (**O** and **P**) Circular correlations between the average Vm-LFP theta phase shift and the average spike phase relative to the LFP theta oscillation for all PV neurons during resting (O, N=28 cells, circular correlation coefficient=0.8042) and during walking (P, N=28 cells, circular correlation coefficient=0.9147). In boxplots, notch indicates median, box indicates 25th to 75th percentiles, whiskers indicate the range of non-outliers, and dots indicate outliers.

Although we recorded only one PV neuron at a time, the tight phase relationship between LFP theta and an individual PV cell’s Vm theta (Figure 4G-J) was conserved across recordings, strongly indicating that Vm theta is synchronized amongst the PV population. Since PV cells play an important role in supporting CA1 LFP theta oscillations, we hypothesized that when individual PV Vm theta and LFP theta are closer in phase, the PV population is more synchronized, leading to greater LFP theta power. Indeed, we found that when Vm-LFP phase shifts of individual PV neuron were close to zero, LFP theta power was significantly higher, suggesting that PV cell synchronization accompanies stronger LFP theta at the network level (Figure 4K and L). Consistent with the lack of synchronization observed between CaMKII Vm and LFP theta, CaMKII Vm-LFP theta phase shifts had no relationship to LFP theta power (Figure 4E and F). Thus, the consistent phase relationship between LFP theta and PV Vm theta, but not CaMKII Vm theta, indicates a unique association of PV neurons with elevated LFP theta power, and supports the prominent role of PV cells in promoting CA1 LFP theta oscillations.

### Transient fluctuations of Vm theta power in both cell types are accompanied by corresponding LFP power changes during resting but not walking

LFP theta oscillation amplitude can be influenced by many sources, including the synchronization of Vm theta between neurons (Figure 4K and L) and the power of Vm theta in individual neurons. To explore how the power of individual neurons’ Vm theta relates to LFP theta power, we aligned LFP theta power to the peak of each Vm theta cycle. During resting, when CaMKII or PV cellular Vm theta power was high (theta cycles with power >0.5), LFP theta and beta power were higher (Supplemental Figure 5A and C, respectively). Similarly, when Vm theta power was low in either cell type (theta cycles with power <-0.5), LFP theta power was correspondingly lower (Supplemental Figure 5B and D, respectively).

Interestingly, during walking, even though we detected an overall increase in LFP theta power (Figure 1), the power of individual Vm theta in both CaMKII and PV cells was no longer associated with transient fluctuations in LFP theta power (Supplemental Figure 5E-H). Thus, transient Vm theta power variations in both pyramidal and PV cells were only accompanied by similar LFP theta power fluctuations when animals were resting but not walking, indicating a behavioral state-dependent coupling between Vm theta power and LFP theta power. These results also suggest that somatic Vm theta power of the CA1 neuronal population is a predominant source of LFP theta only when LFP theta is weak.

### At theta frequency, the PV cell spike-LFP phase relationship captures its underlying Vm-LFP phase relationship, whereas the CaMKII cell spike-LFP phase relationship diverges from its Vm-LFP phase relationship

Since Vm ultimately determines spike timing in individual neurons, one would expect a cell’s spike-LFP temporal relationship to arise from its Vm-LFP temporal relationship. Because extracellular recordings cannot detect Vm, *in vivo* electrophysiology studies are largely limited to observing only the spike-LFP phase relationship during behavior. With our ability to measure Vm, spikes, and LFP simultaneously, we directly examined whether spike-LFP theta phase is a result of Vm-LFP theta coupling. Specifically, we calculated the circular correlation coefficient between the average spike-LFP phase and the average Vm-LFP phase shift of each neuron. As expected, we found that the spike-LFP phases and Vm-LFP phase shifts of individual PV neurons were highly correlated during both resting and walking (Figure 4O and P, circular correlation coefficient, resting: 0.8042, walking: 0.9147). Surprisingly, the spike-LFP phases of individual CaMKII neurons were only loosely correlated with their Vm-LFP phase shifts during both behavioral conditions (Figure 4M and N, N=31 cells, circular correlation coefficient, resting: 0.5313, walking: 0.5257).

We then examined the correlation between spike-LFP phase and Vm-LFP phase shift over a wider range of LFP frequencies and found that these correlations are highest at theta frequency in both CaMKII and PV cells regardless of behavioral condition (Supplemental Figure 6). These results provide direct experimental evidence that the spike-LFP phase relationships of individual PV neurons faithfully represent their underlying Vm-LFP relationships at theta frequency, whereas a CaMKII cell’s Vm-LFP phase relationship is not necessarily revealed by its spike-LFP phase relationship.

## Discussion

LFP oscillations capture rhythmic extracellular potentials and have been broadly linked to behaviors. To understand the cellular mechanisms that support LFP oscillations and organize spike timing, we performed *in vivo* SomArchon voltage imaging of membrane potentials (Vm) from individual CaMKII-positive pyramidal cells or PV-positive interneurons in hippocampal CA1, from mice during resting or walking. We examined the temporal relationships of spikes, Vm oscillations, and simultaneously recorded local field potential (LFP) during various levels of LFP theta oscillations as mice alternated between resting and walking. We present evidence that theta oscillations are a prominent Vm feature in both PV and CaMKII neurons, across behavioral conditions. Vm theta oscillations consistently organize spikes to the rising phase of the Vm theta cycle in both cell types, regardless of the state of LFP theta. Furthermore, spikes and Vm in PV neurons exhibit a tight temporal relationship with CA1 LFP theta phase, supporting the idea that PV cells play a crucial role in promoting hippocampal theta rhythmicity. In contrast, spikes and Vm in CaMKII pyramidal neurons exhibit heterogenous phase relationships with CA1 LFP theta, in line with the idea that CaMKII neurons encode diverse information regarding ongoing behavior. Together, our study provides direct *in vivo* evidence that although cellular theta rhythmicity organizes temporal spike patterns in both pyramidal cells and PV interneurons, PV interneurons play a more prominent role in coordinating CA1 network LFP theta rhythm.

Spike generation requires membrane potential depolarization that reaches action potential threshold. Our results show that spikes preferentially occur on the rising phase of Vm theta, even though the rising and falling phases of a Vm oscillation cycle have similar absolute membrane voltages. This observation confirms that spike threshold is not a fixed value in behaving mice. Instead, transient Vm fluctuations are a critical deterministic criterion for spike generation, as shown in previous work (*39-43*). Specifically, continued monotonic depolarization (rising phase of Vm) leads to significantly greater probability of spike generation at the same absolute membrane voltage than monotonic repolarization (falling phase of Vm).

Additionally, as the large membrane voltage change created by spiking influences voltage-gated ion channels, it is expected that spike occurrence would affect Vm. Indeed, *in vitro* studies have shown that hippocampal pyramidal cell spiking is often followed by an after-depolarization potential that lasts for tens of milliseconds (*47*). We found that in both pyramidal and PV cells, spiking prolongs Vm theta cycles and elevates Vm theta power regardless of behavioral state, highlighting that Vm oscillations and spikes are irrevocably intertwined. One potential biophysical mechanism for this phenomenon is that once Vm reaches action potential threshold, spike generation provides further Vm depolarization and thus amplifies the Vm theta oscillation. Future studies directly measuring ion channel conductance states will help reveal the causal relationships between subthreshold voltage dynamics and suprathreshold spikes.

In accordance with previous studies, we found that spiking in CaMKII pyramidal cells and PV cells occurs at distinct phases of the LFP theta oscillation (*11*). Specifically, CaMKII spikes occurred indiscriminately across the peak, falling phase, and trough of LFP theta, whereas PV spikes were more concentrated to the rising phase of LFP theta. In the hippocampus, silencing or altering PV activity disturbs the timing of pyramidal cell spiking (*6, 7, 15, 21*). Each PV neuron often inhibits multiple pyramidal neurons, and pyramidal cell spiking is strongly influenced by rebound depolarization following PV inhibition (*7, 21*). Thus, our observation that PV spiking often precedes pyramidal cell spiking within an LFP theta cycle suggests that the differences in spike-LFP phase between CaMKII and PV neurons observed here may originate from direct synaptic interactions between these two populations.

Hippocampal LFP theta oscillations capture rhythmic extracellular potential that arises from somatic, dendritic, and axonal Vm fluctuations in individual neurons. These Vm fluctuations are influenced by both rhythmic synaptic inputs to the hippocampus and the intrinsic biophysical properties of individual neurons (*5, 35*). We found that the Vm of both CaMKII and PV cells exhibits strong theta oscillations, regardless of network LFP theta state. Previous intracellular patch clamp studies and voltage imaging studies have reported prominent Vm theta oscillations in CA1 neurons of animals across anesthetized, quiescent, and active locomotion behavioral conditions (*22, 24-27*), even though a recent voltage imaging study failed to detect prominent Vm theta in resting mice (*27*). It is possible that the prominent Vm theta across behavioral states we observed is a result of improved voltage imaging sensitivity via cell type-specific expression of SomArchon, though we cannot rule out other factors such as behavioral state variations. Vm theta oscillations in pyramidal neurons likely originate from both synaptic input and their intrinsic biophysical properties, as they exhibit resonance at theta frequency (*48, 49*). PV neurons, however, exhibit resonance at gamma frequencies (*50*); therefore, Vm theta oscillations in PV neurons are likely driven by rhythmic synaptic inputs, such as those from GABAergic neurons in the medial septum (*16, 19, 20*).

PV cells have been shown to powerfully contribute to CA1 theta oscillations in slice and *in vivo* (*14, 15*). Consistent with these observations, our results show that Vm theta oscillations in PV neurons are better temporally aligned with LFP theta oscillations than Vm theta of CaMKII cells, and higher synchrony between PV Vm theta and LFP theta is associated with higher LFP theta power. Because PV neurons are coupled via gap junctions and they receive theta rhythmic inputs from the GABAergic septum, our observation that PV Vm theta is temporally aligned to LFP theta strongly suggests that Vm oscillations are synchronized across PV neurons (*7, 16-20, 51-53*). Conversely, Vm theta oscillations in CaMKII neurons show diverse temporal relationships with LFP theta oscillations. Given our finding that CaMKII neurons exhibited out-of-phase theta oscillations with each other as a population, it is likely that somatic Vm theta of pyramidal cells contributes minimally to LFP theta oscillations. An alternate, yet compatible, explanation is that the frequencies of CaMKII neurons’ somatic Vm theta oscillations are unstable and therefore result in variable Vm-LFP theta phase shifts. Future studies that simultaneously record multiple CaMKII neurons are necessary to reveal the relationship between the Vm oscillations of individual pyramidal neurons and their combined relationship with bulk LFP oscillations.

In our results, transient cycle-by-cycle Vm theta power of both CaMKII and PV cells is correlated to LFP theta power during resting, indicating that in the absence of strong inputs to the CA1, and thus weak or inconsistent LFP theta, the power of individual Vm oscillations is better associated with CA1 LFP oscillations. Intriguingly, during walking, which induced strong LFP theta oscillations, the power of individual Vm theta cycles of both cell types became decoupled from LFP theta power. During locomotion, the CA1 receives strong synaptic inputs from other brain regions, which are best reflected by changes in the dendritic membrane potential of CA1 neurons (*35, 54-56*). SomArchon is targeted to the cell body and therefore we could only quantify changes in the somatic membrane potential of CA1 neurons, which is likely different from synaptically-driven dendritic membrane potential (*1, 57, 58*). As such, the behavioral state-dependent decoupling of individual Vm cycle power and LFP power that we observed here likely indicates that during movement, LFP theta is dominated by dendritic membrane potential driven by synaptic inputs to the CA1 rather than the somatic Vm theta oscillations of the local CA1 neuronal population. Future studies that directly measure dendritic membrane potentials will help elucidate how CA1 neurons dynamically transform behaviorally-relevant dendritic inputs to defined spiking output patterns.

The observation that spike timing is strongly and consistently phase-locked to Vm theta in both pyramidal and PV neurons predicts that the spike-LFP relationship faithfully reflects the Vm-LFP theta phase relationship in both cell types. In PV neurons, we did indeed observe this expected relationship between spike-LFP theta phase and Vm-LFP theta phase shift. Together with the observation that PV Vm theta is tightly coupled to LFP theta, our results suggest that synchronization of Vm oscillations across the PV population provides each PV cell with a consistent temporal framework that aligns with ongoing LFP theta oscillations. This timing mechanism would allow PV neurons to organize their spike timing relative to LFP theta to pace hippocampal networks. Surprisingly, we only detected this tight temporal relationship between spike-LFP theta phase and Vm-LFP theta phase shift in PV neurons, but not in pyramidal neurons. Thus, the spike-LFP theta phase relationship in pyramidal cells provides little information regarding its underlying Vm-LFP theta phase relationship. One potential explanation is that spike timing in pyramidal neurons is strongly influenced by other factors besides intrinsic Vm theta oscillations, such as rebound depolarization from local PV inhibition or cholinergic inputs from the medial septum (*6, 7*). However, despite their weak relationship, we also found that correlations of spike-LFP phase and Vm-LFP phase shifts in pyramidal neurons were highest in the theta frequency compared to other frequency bands, indicating that Vm theta oscillations are nonetheless more important than other frequencies in influencing pyramidal cell spike timing.

## Acknowledgements

X.H. acknowledges funding from NSF (CBET-1848029, DIOS-2002971) and NIH R01MH122971 R.A.M. acknowledges funding from NIH F31MH123008 and NIH/NIGMS Quantitative Biology and Physiology Fellowship (T32GM008764) through the Boston University Biomedical Engineering Department. E. L. acknowledges funding from Boston University Center for Systems Neuroscience and Kilachand Center. The authors acknowledge the support from the Shared Computing Cluster in Boston University’s Research Computing Services, and the Boston University Micro and Nano Imaging Facility (NIH S10OD024993).

## Author Contributions

H. T., R. A. M. and X. H. designed the experiments. R. A. M conducted all experiments. H. T. and R. A. M. performed data analysis. E. L., H. J. G. and C.C. provided technical assistance. X. H. supervised the study. H. T., R. A. M. and X. H. wrote the manuscript. All authors edited the manuscript.

## Data and materials availability

Data are available from lead contact upon reasonable request.

## Materials and Methods

### Animal Surgery and Recovery

All animal procedures were approved by the Boston University Institutional Animal Care and Use Committee. 13 mice expressing Cre recombinase in parvalbumin-expressing cells (PV-Cre mice, B6;129P2-Pvalbtm1(cre)Arbr/J, JAX stock #017320, Jackson Laboratory) were used. Mice were 8-20 weeks old at the start of experiments. Both male and female mice were used. Animals first underwent surgery to implant a sterilized custom imaging window with an attached guide cannula and LFP electrode that was assembled before surgery. The window assembly consisted of a stainless steel cannula (outer diameter: 3.17 mm, inner diameter: 2.36 mm, height: 1.75 mm, B004TUE45E, AmazonSupply) fitted with a circular coverslip (size 0, diameter: 3 mm, Deckgläser Cover Glasses, Warner Instruments), adhered to the bottom using a UV-curable optical adhesive (Norland Optical Adhesive 60, P/N 6001, Norland Products). The guide cannula (26 gauge, No C315GC-4/SP, PlasticsOne) was fixed at an approximately 60° angle to the imaging cannula and terminated flush with the window surface. The LFP electrode consisted of either an insulated stainless steel wire (diameter: 0.125 mm, No: 005SW-30S, PlasticsOne) soldered to a pin (No: 853-93-100-10-001000, Mill-Max) or a bipolar electrode (wire diameter: 0.125 mm, No: MS303S/3-B-SP, PlasticsOne) and was fixed parallel to the guide cannula, terminating about 200 µm below the window surface.

During surgery, an approximately 3.2mm hole was drilled in the skull (centered at anterior/posterior: -2.0mm, medial/lateral: +1.8mm) and the cortical tissue overlaying the hippocampus was aspirated away to expose the corpus callosum. The corpus callosum was then thinned until the underlying tissue of the CA1 could be visualized through the surgical microscope. The imaging cannula was placed on top of the hippocampus (stratum pyramidal) with LFP electrode located in stratum radiatum, and sealed in place using a surgical silicone adhesive (Kwik-Sil, World Precision Instruments). A hole was drilled posterior to lambda to implant an electrode (No: 853-93-100-10-001000, Mill-Max) for ground reference in LFP recordings. The imaging window and ground electrode were secured in place, and a custom aluminum head-plate was attached to the skull (posterior to the window), using bone adhesive (C&B Metabond, Parkell) and dental cement. All mice were treated with buprenorphine for at least 48 hours after each surgery. Mice were singly-housed after window implantation surgery to prevent damage to the head-plate and imaging window.

After complete recovery from surgery (7+ days), 500-750 nL of virus was infused through the guide cannula into the CA1. Most CaMKII mice (n=4) were infused with AAV9-CaMKII-SomArchon-GFP (titer: 3.2×10^12^ GC/mL, Addgene #126942), and one CaMKII mouse was infused with AAV9-synapsin-SomArchon-GFP (titer: 5.9×10^12^ GC/mL, Addgene #126941). PV mice (n=9) were infused with AAV9-CAG-FLEX-SomArchon-GFP (titer: 6.3×10^12^ – 1.1×10^13^ GC/mL, Addgene #126943) or AAV9-synapsin-FLEX-SomArchon-GFP (titer: 1.28×10^13^ GC/mL). All viruses except AAV9-synapsin-FLEX-SomArchon-GFP were obtained from the University of North Carolina Chapel Hill Vector Core. The plasmid for AAV9-synapsin-FLEX-SomArchon-GFP was designed in-house, created by Epoch Life Science, and packaged into AAV by Vigene Biosciences. Animals were awake and head-fixed during infusion. An internal cannula (33 gauge, No: C315IS-4-SPC, PlasticsOne) was inserted into the guide cannula and infusion was performed using a microinjector pump (UMP3 UltraMicroPump, World Precision Instruments). The internal cannula remained in place for 1 min before infusion. Rate of infusion was 50 nL/min. After infusion, the internal cannula remained in place for 5-10 min before being withdrawn. One PV mouse did not receive a viral infusion, and instead underwent a stereotaxic viral injection surgery prior to window implantation. During surgery, a hole was drilled in the skull targeting the hippocampus (anterior/posterior: -2.0mm, medial/lateral: +1.4mm, dorsal/ventral: -1.6mm from bregma). The injection was performed with a blunt 33-gauge stainless steel needle (NF33BL-2, World Precision Instruments) and a 10 µL microinjection syringe (Nanofil, World Precision Instruments), using a microinjector pump (UltraMicroPump3-4, World Precision Instruments). The needle was lowered over 1 min and remained in place for 1 min before infusion. The rate of infusion was 50 nL/min. After infusion, the needle remained in place for 7-10 min before being withdrawn over 1 min. The skin was then sutured closed with a tissue adhesive (Vetbond, 3M). After complete recovery (7+ days after virus injection), the animal underwent a second surgery for window implantation as described above.

### Animal Habituation

After viral infusion, animals were habituated to experimenter handling and head-fixation on a motorized treadmill. Each animal was habituated to cyclic resting and walking at 11.75 cm/sec (20 seconds each, repeating) on the treadmill 4-5 days a week for at least 3 weeks prior to the start of imaging. Viral expression peaked and remained high 4 weeks after infusion/injection. Imaging was performed about once every seven days and animals were continually habituated in the interim between imaging days.

### Voltage Imaging and LFP Recording

During each imaging session, animals were head-fixed on the treadmill underneath a conventional wide-field microscope equipped with an ORCA-Fusion Digital complementary metal oxide semiconductor (CMOS) Camera (C14440-20UP, Hamamatsu Photonics K.K.) and a 40x NA0.8 CFI APO NIR objective (Nikon). Because each SomArchon molecule has an attached GFP tag that can be used as a static label for SomArchon-expressing cells, the microscope was equipped with a 5 W light emitting diode (M470L4, ThorLabs), an 475/15-nm bandpass emission filter, 525/45-nm bandpass excitation filter, and 495-nm dichroic mirror. To excite and record SomArchon, the microscope was also equipped with a 40-mW 637-nm red laser (Coherent Obis 637-140X), a 706/95-nm band-pass emission filter, and a 635-nm dichroic mirror (Semrock).

Cells were first located using the static GFP tag. SomArchon dynamics were subsequently imaged at approximately 828 Hz (1.2 ms exposure) using HCImage Live (Hamamatsu Photonics K.K.) software. HCImage Live data were stored as DCIMG image files, and further analyzed offline. Each cell was imaged for 5 trials while the treadmill was off and 5 trials while the treadmill was on, for a total of 10 trials. Each imaging trial was 12.07 seconds long, with a break of about 45 seconds between each trial. Resting and running trials were interleaved such that each resting trial was followed by a running trial. After imaging SomArchon dynamics of a cell, each cell was also imaged in the GFP channel for 5 treadmill off trials and 5 treadmill on trials, to identify physiological signals (such as hemodynamics or breathing) that may have contaminated the SomArchon channel. Each GFP trial was 12.07 seconds with a break of about 25 seconds between each trial. The imaging field of view was 122 pixels in height and of variable width.

Local field potential (LFP) was recording simultaneously at 1 kHz during all trials using an OmniPlex system (PLEXON). The camera also sent a TTL pulse to the OmniPlex system at the onset of each imaging trial to synchronize SomArchon recordings and LFP recordings.

### Histology

At the end of the experiments, all mice were transcardially perfused and tissue was processed to confirm SomArchon expression and cannula placement. Mice were perfused with 0.01M phosphate buffered saline (PBS, No: BP2944100, Fisher Scientific) followed by 4% paraformaldehyde (No: 158127, Sigma-Aldrich). Brains were carefully removed and post-fixed in 4% paraformaldehyde for 4-12 hours. Brains were then transferred to 30% sucrose solution for at least 24 hours before sectioning. Brains were sectioned into 50 µm-thick coronal slices using a freezing microtome (SM2010R, Leica). Slices were collected through the entire anterior hippocampus, from at least -1.0mm to -3.0mm relative to bregma. A subset of sections from brains with CaMKII-SomArchon expression were stained with a mouse anti-CaMKIIα/β/γ/δ antibody (sc-5306, Santa Cruz, 1:50) followed by Alexa Fluor 568 goat anti-mouse secondary antibody (No: A-11004, Thermo Fisher Scientific, 1:500). A subset of sections from brains with FLEX-SomArchon expression were stained with a guinea pig anti-PV antibody (GP72, Swant, 1:1000) following by Alexa Fluor 568 goat anti-guinea pig secondary antibody (No: A-11075, Thermo Fisher Scientific, 1:500). All antibodies were diluted in 2% normal goat serum and 0.5% Triton-X (No: T9284, Sigma-Aldrich) in 0.01M PBS. Briefly, sections were first rinsed with 0.01M PBS and a solution of 100mM glycine (No: G7126, Sigma-Aldrich) and 0.5% Triton-X in 0.01M PBS, followed by a 2-hour incubation in blocking buffer containing 5% normal goat serum and 0.5% Triton-X in 0.01M PBS. Sections were then incubated for 24 hours with primary antibody, rinsed with 100mM glycine and 0.5% Triton-X in 0.01M PBS, and incubated with secondary antibody for 2 hours. Slices were then incubated for 10 min with Hoechst 33342 (No: 62249, Thermo Fisher Scientific, 1:10,000 in 0.01M PBS), rinsed with 100mM glycine and 0.5% Triton-X in 0.01M PBS, and rinsed again in 100mM glycine in 0.01M PBS. Slices were mounted on slides (Fisherbrand Superfrost Plus, No: 12-5550-15, Fisher Scientific) using anti-fade mounting medium (ProLong Diamond, No: P36965, Thermo Fisher Scientific). Images were taken on an Olympus FV3000 scanning confocal microscope using a 20× objective.

### Data Analysis

#### Motion Correction

All videos were motion corrected with a custom Python script. To assist motion correction, we first pre-processed each frame to enhance the image. To avoid camera artifacts that occasionally occur at the edges of the image, the edge areas corresponding to 10% of the image width/height were discarded. The image was then high-pass filtered (gaussian filter, sigma=50) to remove any potential uneven background. We further enhance the boundary of high intensity areas by adding 100 times the difference between two low-pass filtered images (gaussian filter, sigma=2 and 1) to the low-pass filtered image (gaussian filter, sigma=1). After pre-processing the images, we performed the motion correction as following. We calculated the cross-correlation coefficient between the processed image and a reference image, and obtained the displacement between the center of the image and the maximum coefficient. we then shifted the original unprocessed image accordingly. For the first trial, we used the mean intensity image of the video as the initial reference image to motion-correct the first trial, and then we used the mean intensity image of the motion-corrected first trial as the new reference image to motion-correct all trials.

#### ROI Selection, Trace Extraction, and Spike identification

A region of interest (ROI) was identified for each cell using the first or second video recorded for that cell. A mean projection image was generated for the video and an ROI was drawn manually using ImageJ. An average fluorescence trace for the ROI was then extracted for each video in MATLAB, and detrended with a double-exponential fitting line (MATLAB fit function, ‘exp2’) or linear detrend (MATLAB fit function, ‘detrend’) to remove the influence of photobleaching.

To identify the spikes, we first high-pass filtered the trace at 5Hz to remove any slow changes (Tf), and generated a moving-window averaged trace (Tm, window length = 21 data points). We then generated the upper trace (Tu), which includes the potential spike:

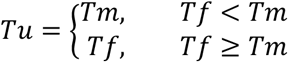

Similarly, we also generated the lower trace (Tl):

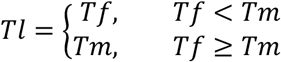

We then calculated the derivatives of Tu and Tl (diff_Tu and diff_Tl, respectively). Since diff_Tl captured the half of the intensity changes not due to spikes, we estimated baseline fluctuation (B) as two times the standard deviation of diff_Tl. Because a spike was a rapid increase in intensity followed by rapid decrease in intensity within a few data points, we identified the data point (t) as a spike with the following two criteria:

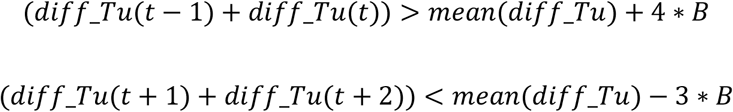

We obtained the amplitude of each spike (A) by calculating the difference between the spike peak intensity and the lower intensity of the prior two data points. We then estimated the corresponding amplitude to baseline fluctuation ratio (ABR). Because true spikes should share a similar ABR, we further refined our spike identification results based on the ABR distribution. Specifically, we performed k-mean clustering on the ABR to obtain two clusters and found the major cluster, which included more spikes. We then excluded the spikes with ABR less than four standard deviations away from the mean of the ABR of the major cluster.

To ensure the quality of the trace, we only included the trials with firing rate > 0.5Hz and average ABR > 1.5, and without motion artifacts identified by manual inspection, blind from experiment conditon. Only ROIs with at least one resting trace and one walking trace were included in further analysis.

### Spectrogram and Power Analysis

Spectrograms were generated with the MATLAB cwt function (‘TimeBandwidth’=60, ‘VoicesPerOctave’=10). We focused all oscillation analysis within the frequency range from 2 to 40Hz, except Supplemental Figure 2. In Supplemental Figure 2, Vm spectrogram was calculated up to 70 Hz to avoid the interference of camera fan mechanical noise around 80Hz, and LFP spectrogram was calculated up to 200Hz. The spectrograms were normalized by z-scoring either over frequencies (to emphasize changes in the power distribution over frequencies; Figure 1 and Supplemental Figure 2) or over time (to emphasize changes in the power of specific frequencies over time; all other figures). To obtain the power of a specific frequency band, the powers within the desired frequency range were averaged. Similarly, to obtain the power of a specific time window, we averaged the powers of all time points within the window. All spectrograms/power distributions from the trials of the same condition (resting/running) from one neuron were averaged to create the representative spectrogram/power of that neuron during the condition, and then averaged across all neurons.

### Bootstrapping Analysis for Significant Changes in Power Distribution

When aligning power distributions to the spike onsets or to the peaks of Vm theta cycles, we obtained the frequency of significant power changes by identifying the frequencies where the population power distribution was outside of the non-specific population power distribution, calculated as following. For true spikes/peaks, we established the potential range of the population power distribution by calculating the average ± standard error across the representative power distributions of all neurons. The non-specific population power distribution was obtained via bootstrapping. Specifically, we randomly selected the same number of timepoints as the spikes/peaks in each session as pseudo-spikes/peaks and performed the same analysis as described in the previous sentence to get a representative non-specific power distribution for a neuron. The representative non-specific power distributions from all neurons were averaged to obtain a non-specific population power distribution. This random selection procedure was repeated 500 times to obtain a set of 500 non-specific population power distributions. We then established the confidence interval of the non-specific population power distribution by calculating the average ± 2*standard deviation of the set of non-specific population power distributions.

### Theta Peak and Cycle Identification and Phase Calculations

To identify theta peaks, we first filtered Vm or LFP at theta frequency (5-10Hz). The theta peaks were defined as the local maximum found by the MATLAB findpeaks function. For each theta peak, the beginning of its corresponding theta cycle was defined as the time point of the minimum intensity between the current peak and the previous theta peak. The end of its corresponding theta cycle was defined as the time point of the minimum intensity between the current peak and the subsequent theta peak. Each cycle was thus defined as the time window between the beginning and end of the cycle. For each cycle, we calculated its actual frequency by taking the inverse of the cycle duration. We calculated the representative power of each cycle by averaging the power across all time points within the cycle.

To obtain the phase of a spike, we calculated a delay-to-period ratio. The delay was defined as the time duration from the peak before the spike to the spike, and the period was defined as the time duration between the two neighboring peaks on either side of the spike. We then converted this ratio to a phase between -π and π, where 0 was the peaks of one oscillation cycle. To then obtain the distribution of spike phase for each neuron, we divided one cycle into 18 bins, and calculated the percentage of the spikes from each neuron that occur within each bin.

When calculating the phase shift between a Vm cycle and LFP cycle, we similarly calculated a delay-to-period ratio and converted the ratio to phase between -π and π. We defined the delay as the time duration from the LFP peak before the Vm peak to the Vm peak. The period was defined as the duration between the two neighboring LFP peaks on either side of the Vm peak. We obtained the distribution of Vm-LFP phase shift for each neuron as described above. Briefly, we divided one cycle into 18 bins and calculated the percentage of phase shifts from each neuron that fell within each bin. To obtain the power of each Vm or LFP cycle, we first calculated the normalized power as described in section Spectrogram and Power Analysis, and used the power at the peak of the oscillation as the power for that cycle. When calculating the LFP power during each Vm oscillation cycle (Figure 4E, F, K, and L), we used the power at the LFP peak closest to the Vm peak. Circular correlation coefficient was calculated with CircStat (*59*).

**Supplemental Figure 1.**
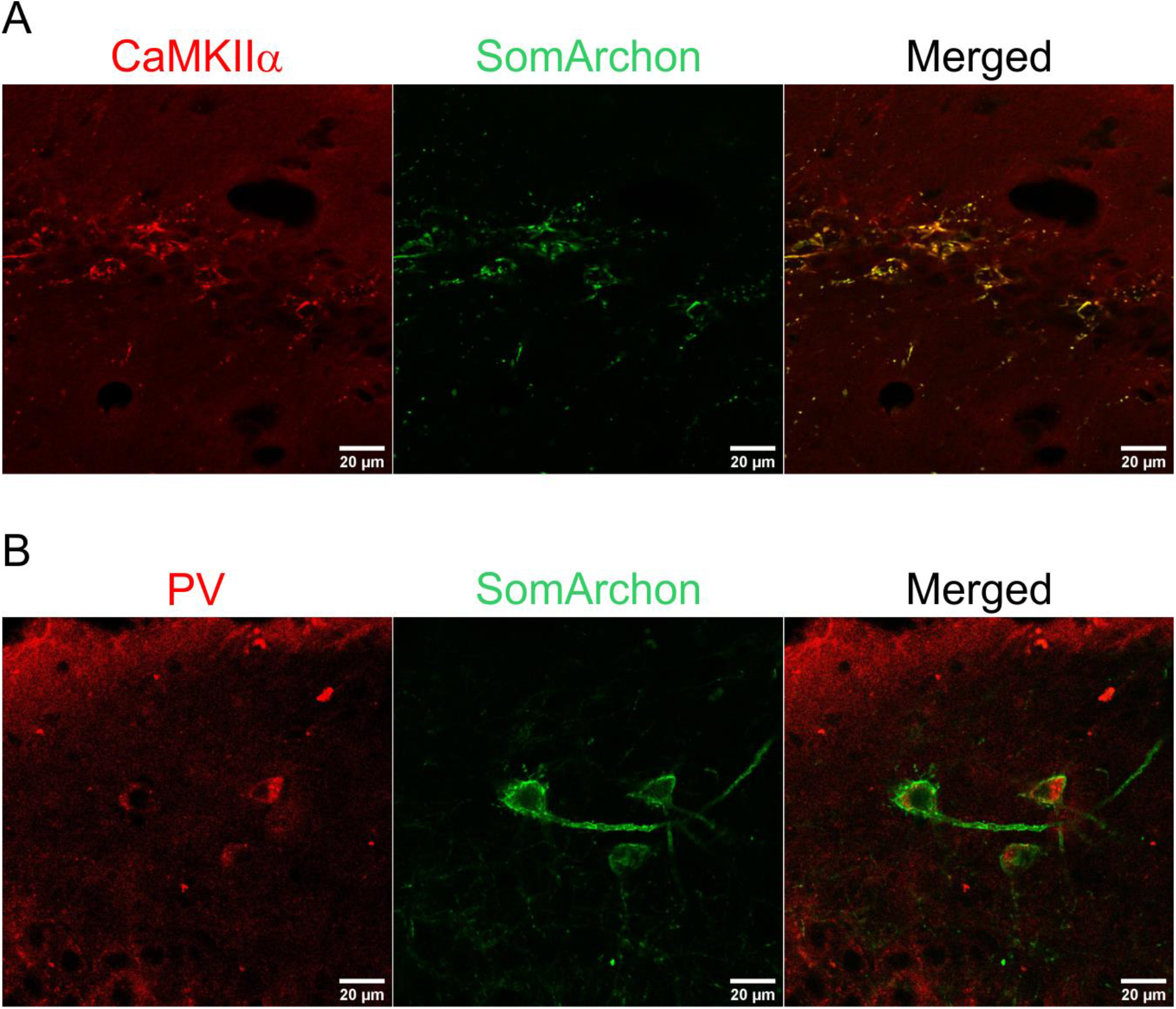
Cell-type specific expression of SomArchon. (**A**) Confocal microscopy images from an example mouse expressing SomArchon in CaMKII cells. Co-localization (right) of CaMKIIα/β/γ/δ immunostaining (left) and CaMKII-SomArchon expression (middle). (**B**) Confocal microscopy images from an example mouse expressing SomArchon in PV cells. Co-localization (right) of PV immunostaining (left) and FLEX-SomArchon expression (middle).

**Supplemental Figure 2.**
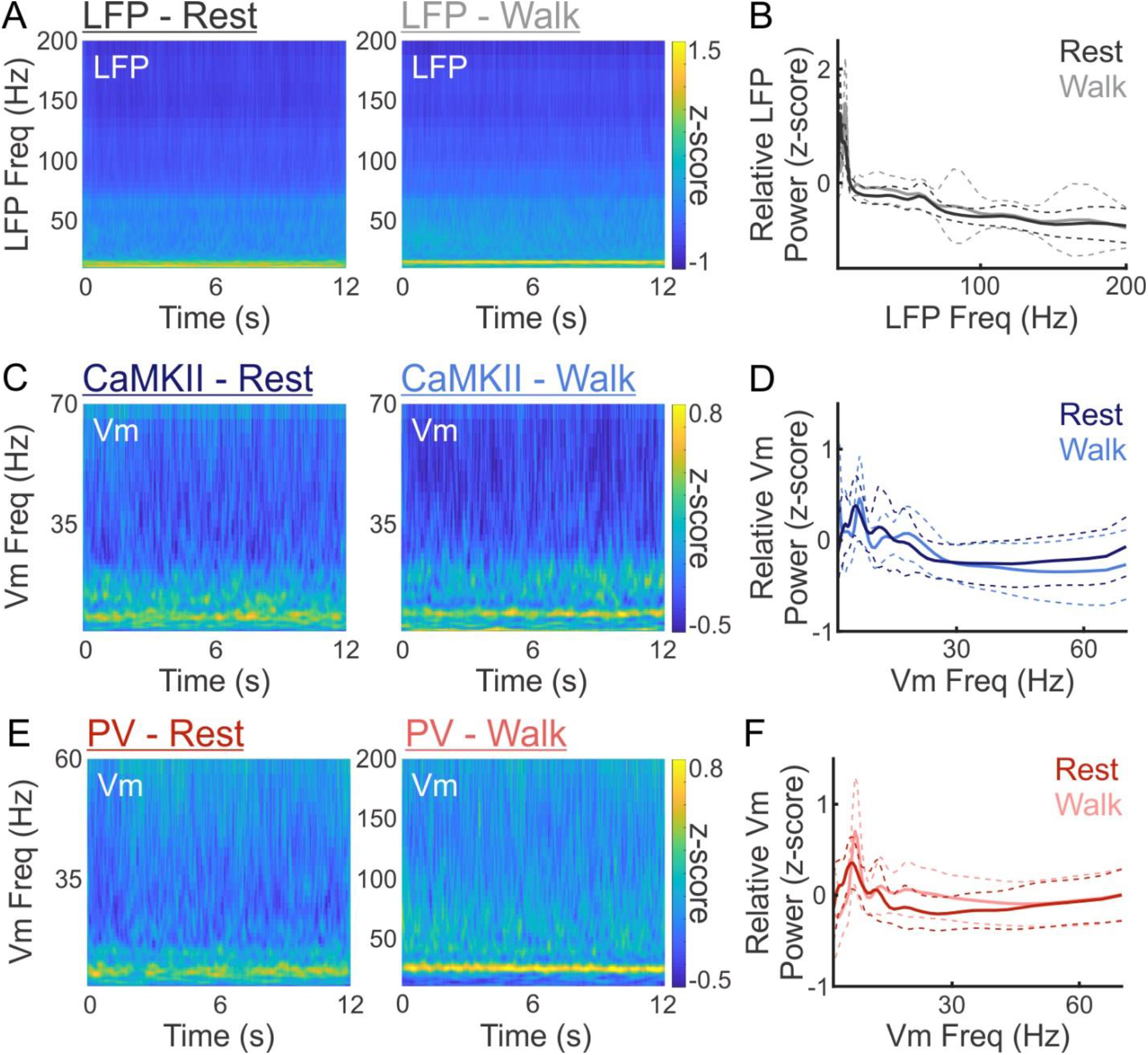
LFP and Vm spectrograms and power during resting and walking. (**A**) Average LFP spectrograms from all recordings, showing a higher frequency range than that shown in Fig 1B, during resting (left) and walking (right) (N= 59 cells). (**B**) Relative LFP power distributions of all recordings during resting (black) and walking (gray) (N= 59 cells). (**C**) Average Vm spectrograms from all CaMKII neurons, showing a higher frequency range than that shown in Fig 1G, during resting (left) and walking (right) (N= 31 cells). (**D**) Relative Vm power distributions of all CaMKII neurons during resting (dark blue) and walking (light blue) (N= 31 cells). (**E**) Average Vm spectrograms from all PV neurons, showing a higher frequency range than that shown in Fig 1K, during resting (left) and walking (right) (N=48 cells). (**F**) Relative Vm power distributions of all PV neurons during resting (dark red) and walking (light red) (N=48 cells). In power plots, solid lines and dashed lines indicate mean and ±standard deviation, respectively. The power at each time point was normalized by calculating its z-score across frequencies. In boxplots, notch indicates median, box indicates 25th to 75th percentiles, whiskers indicate the range of non-outliers, and dots indicate outliers.

**Supplemental Figure 3.**
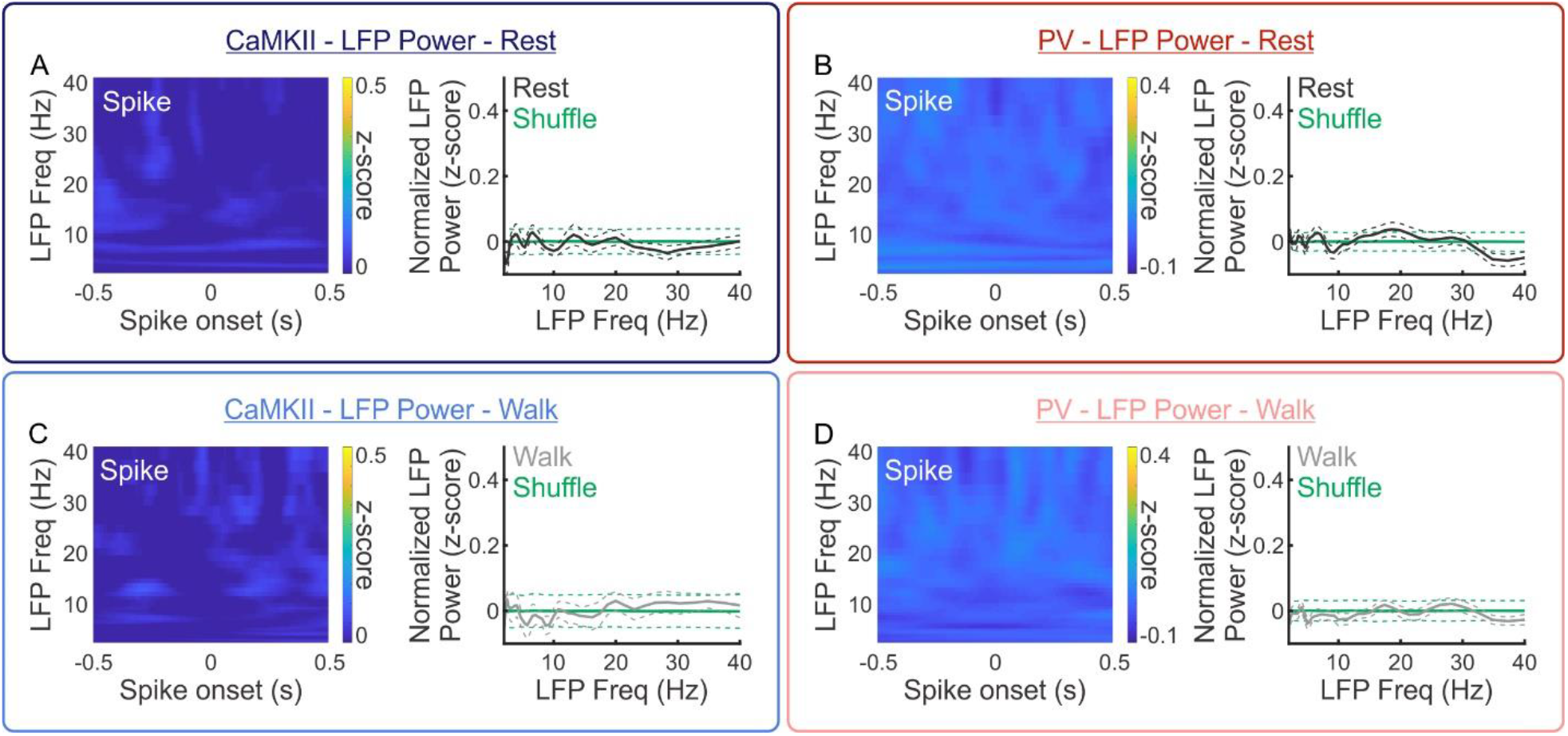
LFP oscillation at spike onsets. (**A**) Average LFP spectrogram (left, N=31 cells) aligned to all CaMKII spikes during resting, and the corresponding LFP power distribution (right, black, mean±standard error, N=31 cells), compared to LFP power in random shuffles (right, green, mean±2*standard deviation). (**B**) Average LFP spectrogram (left, N=28 cells) aligned to all PV spikes during resting, and the corresponding LFP power distribution (right, black, mean±standard error, N=28 cells), compared to LFP power in random shuffles (right, green, mean±2*standard deviation). (**C** and **D**) Same as (A, and B) but during walking (gray). In power plots, the power at each frequency was normalized by calculating the z-score across all time points within each trial.

**Supplemental Figure 4.**
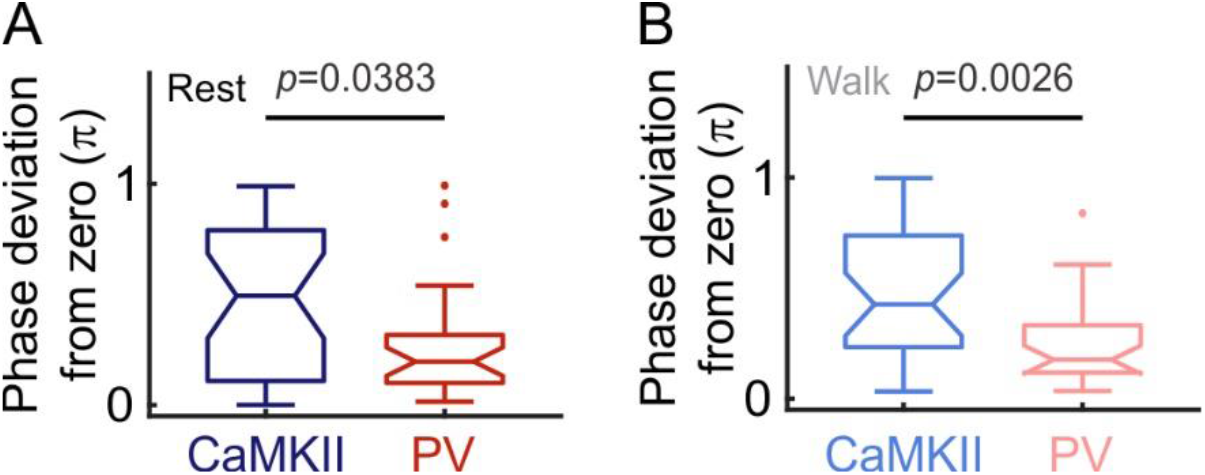
PV Vm theta oscillation is more synchronized with LFP theta oscillation than CaMKII Vm theta oscillation. (**A**) Phase shift between Vm theta oscillation and LFP theta oscillation across all CaMKII neurons (dark blue) and all PV neurons (dark red) during resting (CaMKII: 31 cells, PV: 28 cells, ranksum test, resting: p=0.0383). (**B**) Phase shift between Vm theta oscillation and LFP theta oscillation across all CaMKII neurons (light blue) and all PV neurons (light red) during walking (CaMKII: 31 cells, PV: 28 cells, ranksum test, resting: p=0.0026). In boxplots, notch indicates median, box indicates 25th to 75th percentiles, whiskers indicate the range of non-outliers, and dots indicate outliers.

**Supplemental Figure 5.**
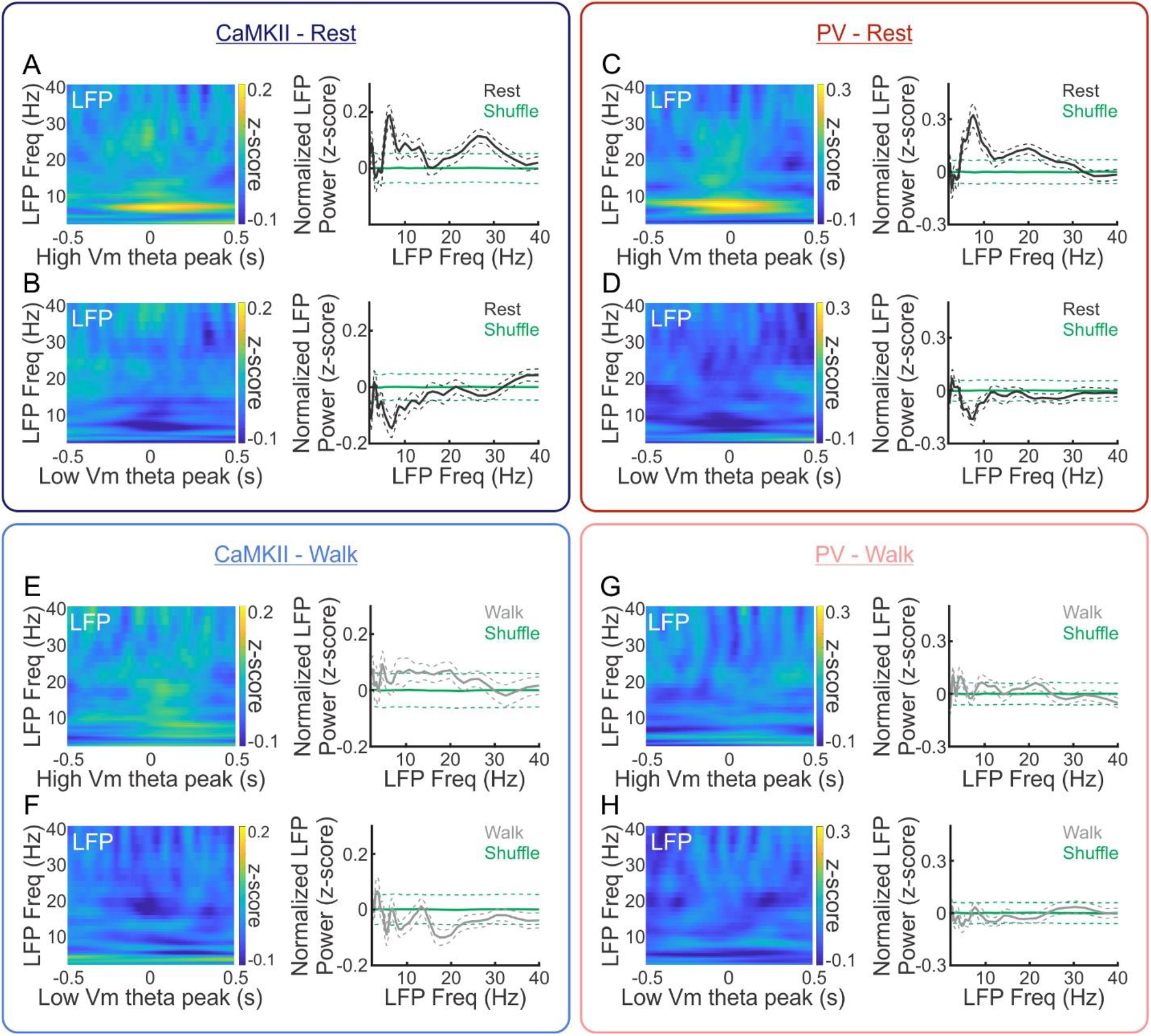
Behavioral state-dependent association between Vm theta power and LFP theta power. (**A**) Average LFP spectrogram aligned to the peak of CaMKII Vm theta cycles with high Vm theta power (power>0.5) during resting (left, N=31 cells), and the corresponding LFP power distribution (right, black, mean±standard error, N=31 cells), compared to the LFP power in random shuffles (right, green, mean±2*standard deviation). (**B**) Average LFP spectrogram aligned to the peak of CaMKII Vm theta cycles with low Vm theta power (power<-0.5) during resting (left, N=31 cells), and the corresponding LFP power distribution (right, black, mean±standard error, N=31 cells) compared to the LFP power in random shuffles (right, green, mean±2*standard deviation). (**C**) Average LFP spectrogram aligned to the peak of PV Vm theta cycles with high normalized Vm theta power (power>0.5) during resting (left, N=28 cells), and the corresponding LFP power distribution (right, black, mean±standard error, N=28 cells) compared to the LFP power in random shuffles (right, green, mean±2*standard deviation). (**D**) Average LFP spectrogram aligned to the peak of PV Vm theta cycles with low normalized Vm theta power (power<-0.5) during resting (left, N=28 cells), and the corresponding LFP power distribution (right, black, mean±standard error, N=28 cells) compared to the LFP power in random shuffles (right, green, mean±2*standard deviation). (**E, F, G**, and **H**) Same as (A, B, C, D), respectively, during walking (gray). In power plots, the power at each frequency was normalized by calculating the z-score across all time points within each trial.

**Supplemental Figure 6.**
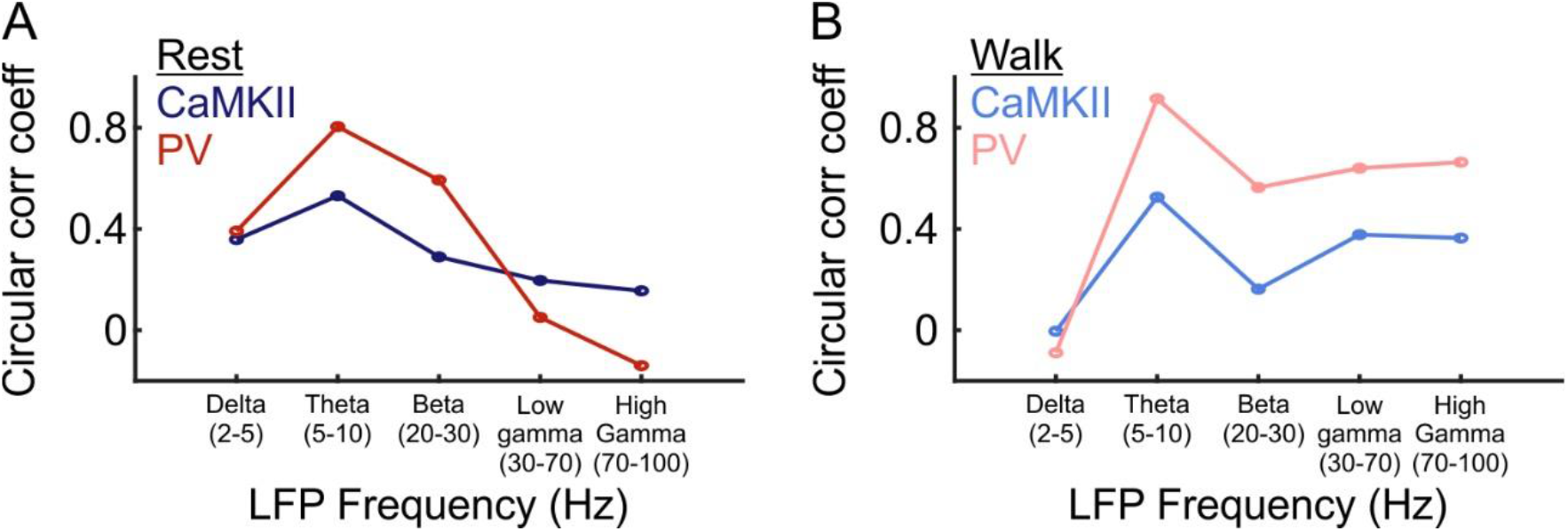
Selective elevation of correlation between spike-LFP phase and Vm-LFP phase shift at theta frequency. (**A**) Circular correlation coefficients between spike-LFP phase and Vm-LFP phase shift across all CaMKII neurons (dark blue) and all PV neurons (dark red) over a range of LFP frequency bands during resting. (**B**) Circular correlation coefficients between spike-LFP phase and Vm-LFP phase shift across all CaMKII neurons (light blue) and all PV neurons (light red) over a range of LFP frequency bands during walking.

